# TSNR: Protein-Specific Signals with Broad Molecular Context for Spatial Protein Prediction

**DOI:** 10.64898/2026.10.27.762369

**Authors:** Xuanyi Zhang, Jingxiao Zhang

## Abstract

Spatial transcriptomics measures transcriptome-wide gene expression while preserving tissue organization, enabling prediction of spatial protein abundance beyond directly measured antibody panels. Such prediction draws on protein-specific transcript signals at the center spot and broader molecular context from the surrounding tissue. Existing spatial methods increasingly use context, but what molecular information the context should retain and how reliably it transfers across tissue environments remain unclear. We compare predictors that share a protein-specific center representation but differ in contextual molecular scope. Their transferred risk difference admits a value-mismatch characterization: broader context is favored when its target-side value advantage exceeds the corresponding change in source-to-target transfer mismatch. For convex prediction objectives, we further relate mismatch to cross-environment estimating-score variation; squared loss yields an explicit covariance geometry. This analysis motivates broad molecular context together with bounded-score robust regression. We develop Target-Specific Niche Regression (TSNR), which combines a protein-specific center representation, compact broad molecular context, and a protein-specific Huber regressor. On the six-section Spatial-CITE-seq benchmark, TSNR achieves the highest mean macro Spearman correlation among the reproduced methods under matched biological holdout evaluation. Controlled analyses support both broader context and robust regression. Across two independent external cohorts, protein-level broad-context gains are positively rank-correlated with the value-mismatch balance in all 11 held-out biological folds.

## 1 Introduction

Spatial transcriptomics measures transcriptome-wide gene expression while preserving tissue organization (Ståhl et al., 2016; Moses and Pachter, 2022; Moffitt et al., 2022). Protein abundance provides complementary information because RNA–protein relationships are shaped by translation, post-transcriptional regulation, and degradation (Liu et al., 2016; Buccitelli and Selbach, 2020). Yet paired spatial protein measurements remain constrained by antibody panels and multiplexing capacity (Liu et al., 2023), motivating prediction of protein abundance from spatial transcriptomics.

Two information sources are especially relevant: protein-specific transcript signals at the center spot and molecular state in the surrounding tissue. RNA-to-protein models establish the predictive value of transcriptomic signals (Zhou et al., 2020; Lakkis et al., 2022; Hanhart et al., 2024), while recent spatial methods incorporate neighborhoods through spatial features, graphs, or context-guided inference (Wang et al., 2026a,b; Lu et al., 2027). However, they do not directly examine whether contextual molecular scope should inherit the protein-specific restriction used at the center when prediction transfers across biological environments.

The center and its spatial context play different representational roles, so their molecular scope need not be symmetric. At the center, protein-specific selection focuses the representation on transcripts associated with the target protein. Applying the same restriction to neighboring spots, however, may discard broader tissue-state signals that are only indirectly related to the target but remain informative for its spatial abundance. Retaining a wider contextual scope can recover such information, but it can also introduce predictive mappings that vary more strongly across tissues. We therefore study how broad the molecular scope of spatial context should be for prediction across biological environments.

We characterize this trade-off at the level of the complete predictor. Comparing target-restricted and broader context yields a *value-mismatch* characterization in which transferred advantage reflects the change in target-side predictive value relative to the change in source-to-target mismatch. For convex objectives, mismatch is further related to cross-environment estimating-score variation; squared loss gives an explicit covariance-weighted geometry.

These results motivate two design choices: broader contextual molecular scope when its value offsets transfer mismatch, and bounded-score robust regression to limit residual-driven score variation. Target-Specific Niche Regression (TSNR) implements these principles with a protein-specific center representation, a compact broad molecular niche shared across targets, and a separate Huber regressor for each protein.

On the six-section Spatial-CITE-seq benchmark, TSNR reaches a mean macro Spearman correlation of 0.4391 versus 0.4164 for the strongest reproduced baseline. Controlled analyses support broader context and robust regression. Across two independent external cohorts, protein-level broad-context gains are positively rank-correlated with the value–mismatch balance in all 11 held-out biological folds.

Our main contributions are:

- We characterize broad-context transfer through a predictor-level value–mismatch decomposition, together with a score-shift bound for convex objectives and a closed-form squared-risk geometry;
- We develop TSNR, combining protein-specific center signals, broader molecular context, and bounded-score Huber regression; and
- We provide protocol-matched experiments showing improved spatial protein prediction, controlled support for both design choices, and external-cohort alignment between value– mismatch ordering and contextual transfer.

## 2 Related Work

### 2.1 Protein-Specific Signals for RNA-to-Protein Prediction

RNA-to-protein prediction spans nonlinear neural networks, simpler regression models, and receptor-specific transcript selection (Zhou et al., 2020; Lakkis et al., 2022; Hanhart et al., 2024; Javaid and Frost, 2023). These studies show that informative transcripts vary across protein targets, but do not address how protein-specific center signals should be combined with molecular context from surrounding tissue.

### 2.2 Spatial Context Representations

Spatial protein-prediction methods incorporate local context through spatial features, graph-based representations, or context-guided inference, including SR2P, DGAT, and SPINE (Wang et al., 2026a,b; Lu et al., 2027). More broadly, spatial-omics methods such as COVET, ENVI, and NicheCompass represent tissue state through neighborhood-level molecular structure or spatial gene programs (Haviv et al., 2025; Birk et al., 2025). However, whether protein-prediction context should share the center’s target-specific molecular restriction remains unclear. TSNR focuses on the molecular scope of the contextual representation.

### 2.3 Generalization Across Environments

Domain adaptation and domain generalization address prediction under distribution shift through distribution discrepancy, invariant representations, and related strategies (Ben-David et al., 2010; Muandet et al., 2013). We consider a more specific transfer problem: after accounting for a protein-specific center representation, the contribution of context depends on both the additional predictive value it retains and the mismatch of its predictive mapping across environments. The value-mismatch characterization formalizes this representation-level transfer trade-off.

## 3 Methodology

### 3.1 Overview

Figure 1 summarizes the representation principle underlying TSNR and its relation to the value– mismatch characterization.

**Figure 1:**
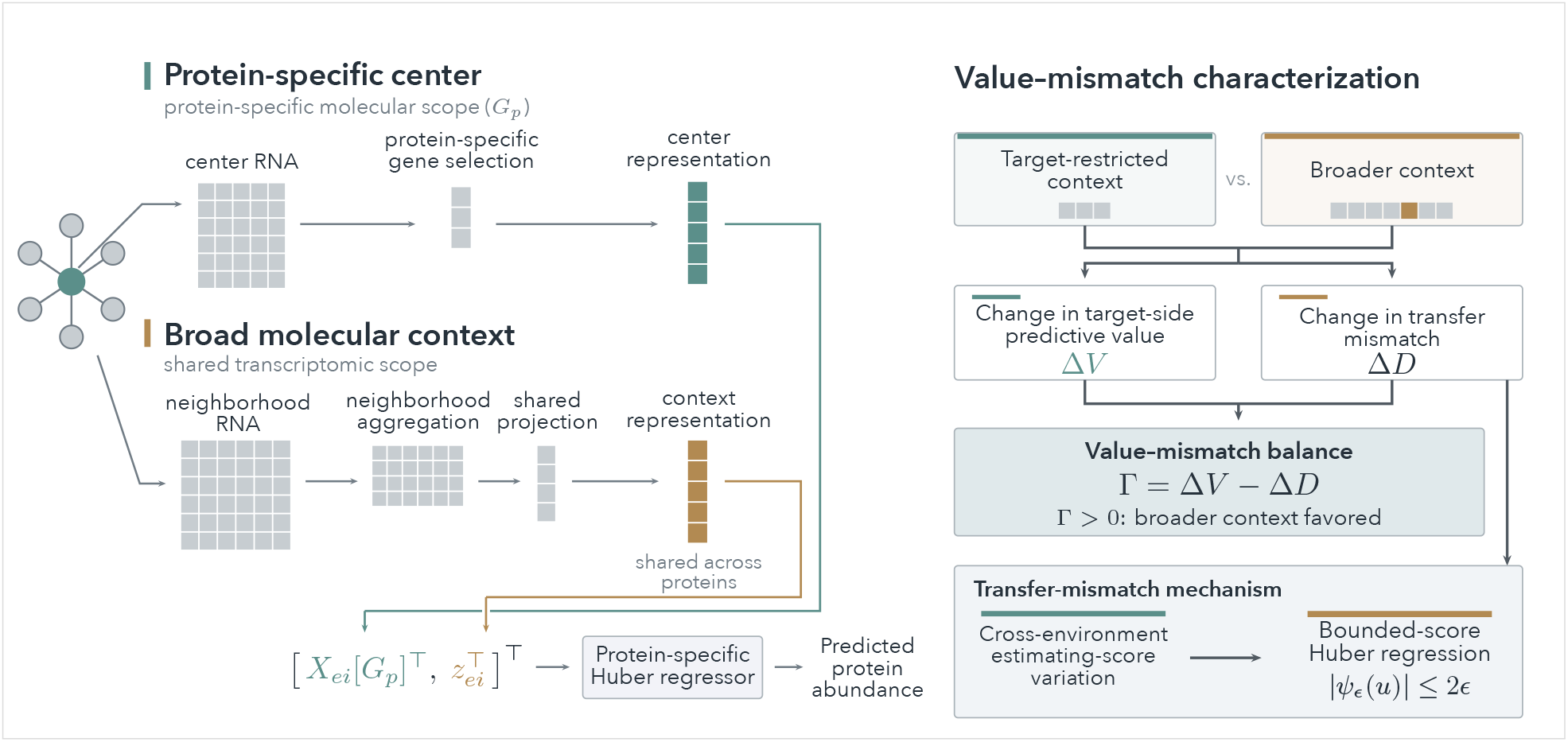
Overview of TSNR and the value–mismatch characterization. **Left:** TSNR combines a protein-specific center representation with broader molecular context and a protein-specific Huber regressor. **Right:** broad and target-restricted predictors are compared through Γ = Δ*V* − Δ*D*, balancing target-side predictive value against transfer mismatch. Cross-environment estimating-score variation is related to transfer mismatch and motivates the bounded-score Huber regression used by TSNR.

We formulate transfer at the level of the complete signal–context predictor. The value–mismatch characterization separates target-side value from transfer mismatch and relates the latter to cross-environment estimating-score variation, motivating broad molecular context together with robust regression.

### 3.2 Signal–Context Prediction Setup

Let *e* ∈ *ℰ* index a tissue environment, *i* ∈ *ℐ*_*e*_ a spatial location, and *p* ∈ *P* a protein target. Let *N*_*ei*_ ⊆ *ℐ*_*e*_ denote the spatial-neighbor index set. We denote center transcriptomics by *X*_*ei*_ ∈ ℝ^*G*^ and protein abundance by *Y*_*eip*_. For protein *p*, define

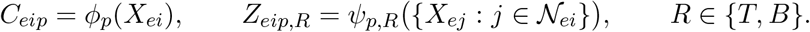

where *T* denotes target-restricted context and *B* broader molecular context; here “target-restricted” refers to the protein target rather than the target environment. The complete representation is

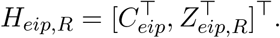

In our evaluations, an environment corresponds to a complete biological holdout unit: a tissue section for Spatial-CITE-seq, a patient for GSE289326, and a specimen for CytAssist.

Let *t* be the held-out target environment and *s* the pooled source environments. After source-side representation fitting, the maps are fixed. For *q* ∈ {*s, t*}, let *J*_*q,p,R*_(*η*) denote the complete prediction objective, including estimator regularization when present, and

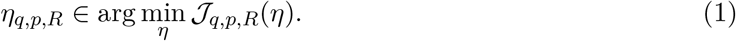

We use *η*_*t,p,R*_ as the target optimum under the fixed source-derived representation.

### 3.3 Value-Mismatch Characterization

We compare the source-trained predictors using broader and target-restricted context directly on the target environment. Define their transferred broad-context advantage as

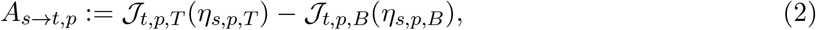

so that *A*_*s*→*t,p*_ *>* 0 indicates lower target objective for the source-trained broad-context predictor.

The target-side value difference between the two representations is

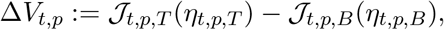

while the source-to-target transfer mismatch within regime *R* is

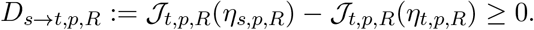

Let

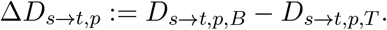

A direct rearrangement gives

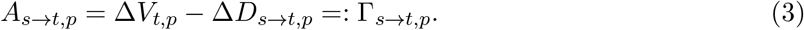

Thus, broader context transfers favorably when its target-side value advantage exceeds the corresponding change in transfer mismatch. Because the two contextual regimes need not be nested, both differences may have either sign.

#### Transfer-mismatch control

The value–mismatch identity isolates transfer mismatch as the term that can offset the predictive value retained by broader context. This mismatch can itself be related to the behavior of the prediction objective across environments. For a fixed regime *R*, suppose that *J*_*t,p,R*_ is differentiable and convex in *η*. By first-order convexity at the source solution,

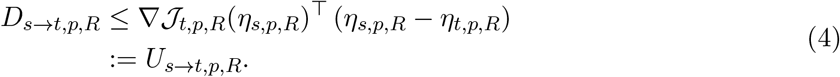

If the source solution is an interior stationary point, then ∇*J*_*s,p,R*_(*η*_*s,p,R*_) = 0, and the target score can equivalently be written as the cross-environment score shift

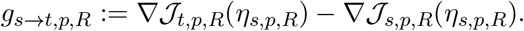

Hence transfer mismatch is controlled by the interaction between cross-environment score variation and the displacement between the source- and target-optimal predictors. Combining this result with Eq. 3 also yields lower bounds and sufficient conditions for favorable broad-context transfer; these refinements are given in Appendix A.

#### Squared-risk geometry

For a linear predictor under squared loss, augment *H*_*p,R*_ with an intercept and define 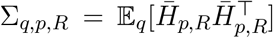 and 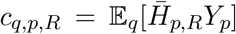. The canonical optimum is 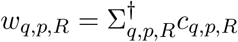, and the target mismatch becomes

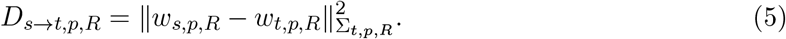

Thus squared risk gives an explicit covariance geometry for the same transfer-mismatch quantity. The corresponding value expression, score-shift form, and singular case are derived in Appendix A.

### 3.4 Target-Specific Niche Regression

The analysis above yields two design implications. First, given a protein-specific center representation, the contextual branch need not inherit the same molecular restriction when broader context retains enough target-side value relative to its transfer penalty. Second, Eq. 4 motivates limiting the sensitivity of the estimating score to occasional large residuals. TSNR provides a simple realization of these two principles through a protein-specific center representation, a compact broad molecular niche, and protein-specific robust regression.

#### Protein-specific center representation

For a given outer split, let *S* denote the set of training environments. For each protein *p*, genes are ranked according to RNA–protein associations estimated from these environments. Let 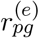 denote the Pearson correlation between gene *g* and protein *p* in training environment *e* ∈ *S*. We define

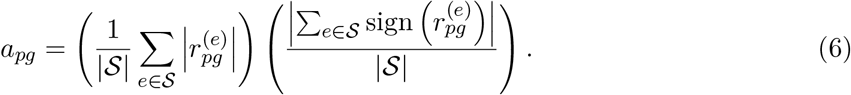

The score favors genes with strong and directionally consistent associations across training environments. The top *K* genes define *G*_*p*_, yielding

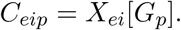

#### Broad molecular niche

For each environment *e*, the spatial neighbor set *N*_*ei*_ is constructed from section-local coordinates. Instead of applying the protein-specific restriction *G*_*p*_, TSNR aggregates the shared transcriptomic space of neighboring spots and projects it into a compact representation,

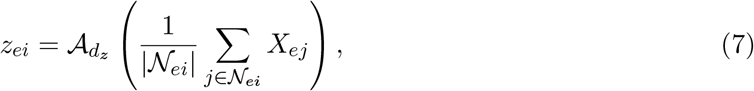

where 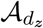 denotes the *d*_*z*_-dimensional PCA transform fitted using neighborhood profiles from the training environments only. In the broad regime, *Z*_*eip,B*_ = *z*_*ei*_ for all proteins, so the contextual representation is shared across protein targets while its predictive contribution remains protein specific.

For comparison, the target-restricted regime uses 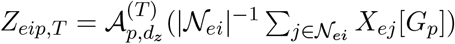, where 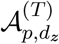 is fitted using the training environments only.

#### Protein-specific robust regression

For each protein, the center and niche representations are concatenated as

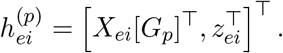

Equation 4 identifies the estimating score at the source solution as a factor controlling transfer mismatch. This motivates controlling the residual-dependent contribution to the estimating score. We therefore use Huber regression (Huber, 1964), whose residual score is bounded. Huber regression provides a simple convex realization of this requirement.

For a standardized residual *u*, define the Huber loss

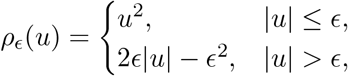

with residual score

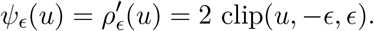

Hence

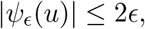

so the residual-dependent factor in the coefficient score remains bounded even for large residuals, while moderate residuals retain the quadratic behavior of squared loss. TSNR therefore fits a separate Huber regressor for each protein. The implementation jointly estimates the regression coefficients, intercept, and residual scale, so the clipping is applied to scale-adaptive standardized residuals. Estimator settings are in Appendix B.2; the finite-sample objective is in Appendix C.1.

The residual behavior motivating robust regression and comparisons with alternative estimators are examined in Sec. 4.3 and Appendix C.

## 4 Experiments

### 4.1 Experimental Setup

#### Datasets

Our primary benchmark comprises six Spatial-CITE-seq sections (Liu et al., 2023) (two tonsil, one breast cancer, one glioblastoma, and two mesothelioma), with 31 protein assay targets. We use six-fold leave-one-section-out evaluation, fitting all fold-dependent transformations on the training sections. For external value–mismatch analysis, we additionally use GSE289326 (Chang et al., 2025), evaluated leave-one-patient-out across six breast cancer patients, and five CytAssist FFPE tumor specimens (Cervilla et al., 2026), evaluated leave-one-specimen-out. Both external cohorts are evaluated on the same 31 assay features, corresponding to 30 unique protein identities.

#### Baselines

We compare TSNR with three recent state-of-the-art spatial RNA-to-protein prediction methods: DGAT (Wang et al., 2026a), SPINE (Lu et al., 2027), and SR2P (Wang et al., 2026b). All methods use the same outer biological splits and evaluation metric, while method-specific modeling and preprocessing are retained as closely as possible. Reproduction details are given in Appendix B.4.

#### Evaluation Metric

For each held-out environment, we compute Spearman correlation between predicted and measured abundance for each of the 31 assay targets and report their macro average, weighting biological holdout folds equally.

#### Implementation Details

TSNR uses *K* = 256 center genes per protein, six non-self neighbors from a 36-nearest-neighbor candidate graph, a *d*_*z*_ = 64 PCA context representation, and Huber regression with robustness parameter *ϵ* = 1.35 and *ℓ*_2_ penalty *α* = 10^−4^. The reported configuration is applied unchanged to the external cohorts without cohort-specific retuning, and all fold-specific transformations are fitted using outer-training data only. Development, preprocessing, implementation, and sensitivity details are provided in Appendix B.1, B.2, and C.4.

### 4.2 Main Results

Table 1 compares TSNR with the reproduced baselines under the common six-fold biological holdout protocol. TSNR achieves the highest mean macro Spearman correlation of 0.4391, compared with 0.4164 for DGAT, 0.3536 for SPINE, and 0.2464 for SR2P. Relative to the strongest reproduced baseline, DGAT, this corresponds to a 5.5% improvement in mean correlation. This improvement is not concentrated in a single tissue or a small subset of assay targets: TSNR outperforms each of DGAT and SPINE on five of the six held-out sections and SR2P on all six, while 65.1% of the 186 protein–fold TSNR–DGAT differences are positive and 20/31 proteins have positive fold-averaged differences. Performance nevertheless varies across tissues, with SPINE slightly stronger on Tonsil-1 and DGAT attaining the highest score on glioblastoma.

**Table 1:** Macro Spearman correlation under six-fold leave-one-section-out evaluation. Best results are bold; the last column reports the relative gain of TSNR over each baseline in mean correlation. GBM denotes glioblastoma and Meso denotes mesothelioma.

| Method | Tonsil-1 | Tonsil-2 | Breast | GBM | Meso-1 | Meso-2 | Mean | TSNR Gain |
|---|---|---|---|---|---|---|---|---|
| SR2P | 0.3269 | 0.4020 | 0.3577 | 0.0969 | 0.1511 | 0.1438 | 0.2464 | +78.2% |
| SPINE | **0.5198** | 0.6142 | 0.4192 | 0.0962 | 0.2281 | 0.2439 | 0.3536 | +24.2% |
| DGAT | 0.5110 | 0.6179 | 0.5111 | **0.2200** | 0.3223 | 0.3159 | 0.4164 | +5.5% |
| **TSNR (ours)** | 0.5142 | **0.6573** | **0.5899** | 0.1971 | **0.3346** | **0.3414** | **0.4391** | – |

#### Qualitative examples

Figure 2 shows two held-out protein–section examples comparing TSNR with DGAT, the strongest baseline. For both CD3E and CCR7, TSNR better preserves the measured spatial structure than DGAT and yields higher protein-level Spearman correlations.

**Figure 2:**
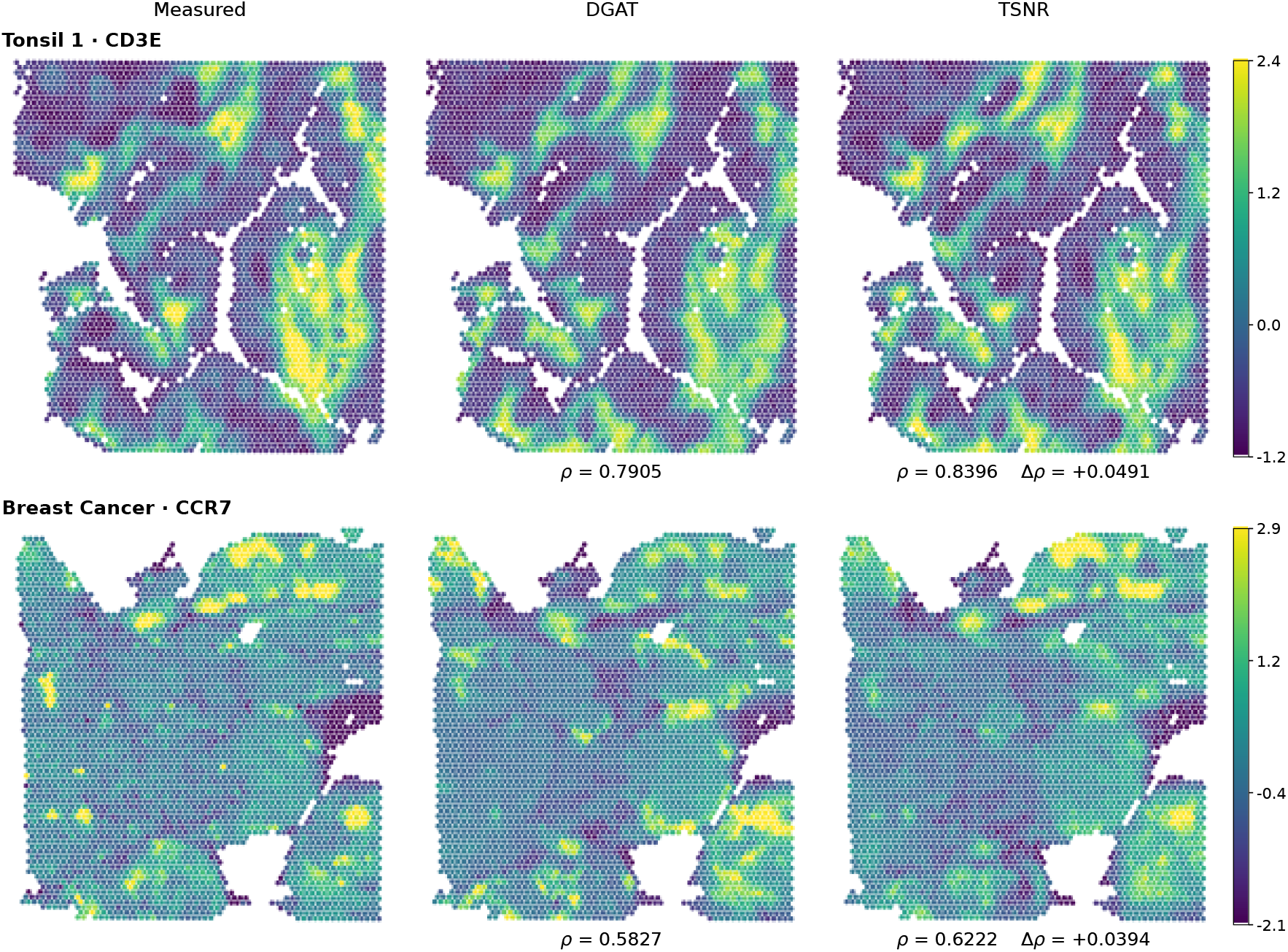
Illustrative held-out spatial predictions from the primary Spatial-CITE-seq benchmark. Panels show measured abundance and predictions from DGAT and TSNR; *ρ* denotes protein-level spatial Spearman correlation and Δ*ρ* = *ρ*_TSNR_ − *ρ*_DGAT_.

### 4.3 Ablation Studies and Analyses

We next examine the two design choices motivated by the analysis in Sec. 3.3: retaining broader molecular information in the contextual representation and using robust protein-specific regression.

#### Molecular scope of context

We first vary the contextual representation while keeping the protein-specific center representation and regression framework fixed. Target-restricted context uses the same protein-specific gene set *G*_*p*_ as the center, whereas TSNR retains the broader transcriptomic neighborhood; both use the same fitted context dimension. We additionally consider a correspondence-disrupted broad context and a size-matched random-gene control. Exact constructions are given in Appendix B.3.

Target-restricted context improves center-only performance (0.4241 vs. 0.4148), while broad context reaches 0.4391. With context dimension fixed, the broad-over-restricted gain remains +0.0131 after excluding *G*_*p*_ (+0.0150 for full broad); neither disrupted correspondence nor random genes reproduces the gain.

#### Robust protein-specific regression

Motivated by the score–mismatch relation in Sec. 3.3, we compare the fixed TSNR Huber estimator with penalty-matched and source-tuned Ridge under the same representation.

Huber has the higher fold-level score in every biological holdout under both comparisons. Against the source-tuned Ridge comparator, the mean fold-level Huber advantage across the 17 holdouts is +0.0341. Residual diagnostics on the primary benchmark are consistent with the robust-regression setting: across 186 out-of-fold protein-by-fold residual groups, the median excess kurtosis is 2.33 and all groups have positive excess kurtosis, while the largest 1% of residuals account for approximately 14.8% of the total squared loss. Under controlled response contamination, Huber remains nearly stable as the contamination fraction increases, whereas Ridge progressively degrades. Ridge tuning, residual diagnostics, and contamination results are given in Appendix C.5.

Additional estimator, representation, and center-gene-set sensitivity analyses are reported in Appendix C.4.

### 4.4 Value–Mismatch Analysis

We next examine whether the value–mismatch characterization tracks protein-level variation in broad-context transfer across biological environments.

For each held-out environment and protein, we evaluate the same Huber objective used by TSNR under the target-restricted and broad representations. Let 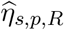 denote the source-trained Huber predictor and 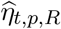 the corresponding target-reference diagnostic fit under the same fixed representation and estimator settings. Using the finite-sample target objective 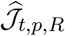, we compute

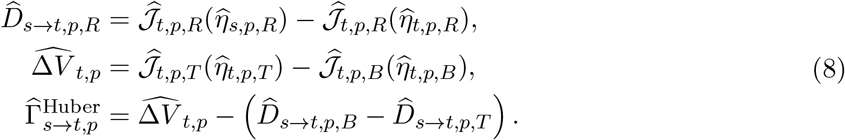

This is the finite-sample Huber counterpart of Eq. 3. Its construction is detailed in Appendix C.1.

We compare 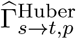 with the empirical protein-level broad-context gain Δ*ρ*_*p*_ = *ρ*_broad,*p*_ − *ρ*_restricted,*p*_ using Spearman correlation across assay targets within each biological holdout. Here, “broad” denotes the TSNR representation and “restricted” denotes the target-restricted matched-neighborhood variant in Table 2. This diagnostic is retrospective and does not enter TSNR model fitting.

**Table 2:** Controlled comparisons of the contextual representation on the six-section Spatial-CITE-seq benchmark. The center representation and regression framework are fixed across all variants. The correspondence-disrupted and random-gene controls are averaged over three and five fixed randomizations, respectively.

| Variant | Context content | Context source | Mean |
|---|---|---|---|
| Center only | None | – | 0.4148 |
| Correspondence-disrupted | Broad transcriptome | Disrupted correspondence | 0.4141 |
| Random-gene control | Size-matched random genes | Matched neighborhood | 0.4164 |
| Target-restricted | Target-restricted | Matched neighborhood | 0.4241 |
| TSNR | Broad transcriptome | Matched neighborhood | **0.4391** |

**Table 3:** Fold-wise comparison of Huber with Ridge using the same TSNR representation. Entries report the number of biological holdouts in which Huber attains the higher macro Spearman correlation.

| Ridge comparator | Spatial-CITE-seq | GSE289326 | CytAssist | All folds |
|---|---|---|---|---|
| α = 10<sup>−4</sup> | 6/6 | 6/6 | 5/5 | 17/17 |
| Source-tuned | 6/6 | 6/6 | 5/5 | 17/17 |

**Table 4:** Within-fold association between the Huber value–mismatch balance and protein-level broad-minus-restricted contextual gains on two independent external cohorts. The middle column reports the mean of the fold-wise Spearman correlations across assay targets.

| Cohort | Mean within-fold Spearman | Positive folds |
|---|---|---|
| GSE289326 | 0.488 | 6/6 |
| CytAssist | 0.478 | 5/5 |
| External total | – | 11/11 |

The association is positive in every external biological holdout. Its magnitude varies across environments, but the protein-level ordering induced by the Huber value–mismatch balance remains aligned with the observed broad-context gain in both cohorts. The corresponding mean association on the six-section primary benchmark is 0.749, with positive correlations in all 6/6 folds.

The component analysis further shows that transfer-mismatch variation is more closely associated with the observed protein-level ordering than target-side value alone. Across all 17 biological holdouts, the mean within-fold Spearman correlations with protein-level contextual gain are 0.108 for 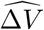, 0.467 for 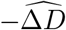, and 0.577 for the complete 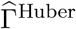. The complete balance is more strongly associated with the observed gain than the value term alone in 16 of 17 folds and than the mismatch term alone in 14 of 17 folds. The Huber-aligned diagnostic also shows a higher mean association with contextual gain than the squared-risk specialization (0.577 versus 0.392) and is stronger in 15 of 17 biological holdouts.

The association also persists when the diagnostic and evaluation outcomes are computed on spatially disjoint target blocks. Across the two directions of this cross-block analysis, 33 of 34 correlations are positive, with a mean of 0.480; after averaging the two directions within each biological holdout, the association is positive in all 17 folds.

#### Empirical score–mismatch alignment

The first-order quantity in Eq. 4 also closely tracks the empirical transfer mismatch of the fitted Huber predictors. Across 1,054 protein–fold–regime fits, the within-analysis Spearman association between 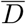 and *U* is approximately 0.98, with median 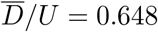. More details are reported in Appendix C.3.

The aggregate effect of broader context nevertheless differs across the two external cohorts: the mean broad-minus-restricted macro Spearman difference is +0.0157 on GSE289326 and −0.0142 on CytAssist. On CytAssist, the normalized target-side value difference is positive in 96% of protein– fold pairs 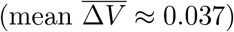, but the larger mismatch increase 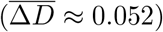 yields 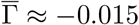. This illustrates how broader context can retain predictive value while transferring unfavorably when the corresponding mismatch increase is larger. Cohort-wise cross-block, squared-risk, and CytAssist component results are reported in Appendix C.2.

### 4.5 Computational Efficiency

Table 5 reports end-to-end wall-clock runtime under each method’s final execution configuration; TSNR requires 126.8 seconds per biological holdout fold on CPU.

**Table 5:**
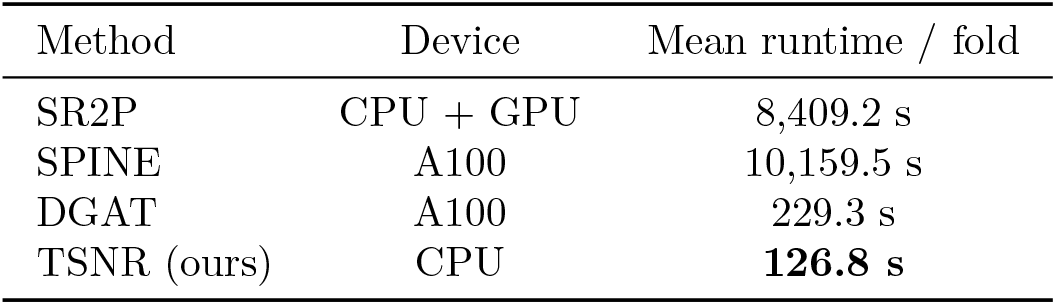
End-to-end wall-clock runtime on the six-section Spatial-CITE-seq benchmark, averaged over six biological holdout folds. Results reflect each method’s final execution configuration.

**Table 6:** Spatially disjoint target-block analysis. Each fold contributes two directional comparisons, one for each assignment of the diagnostic and evaluation blocks.

| Cohort | Positive directions | Directions |
|---|---|---|
| Spatial-CITE-seq | 12/12 | 12 |
| GSE289326 | 12/12 | 12 |
| CytAssist | 9/10 | 10 |
| All cohorts | 33/34 | 34 |

**Table 7:**
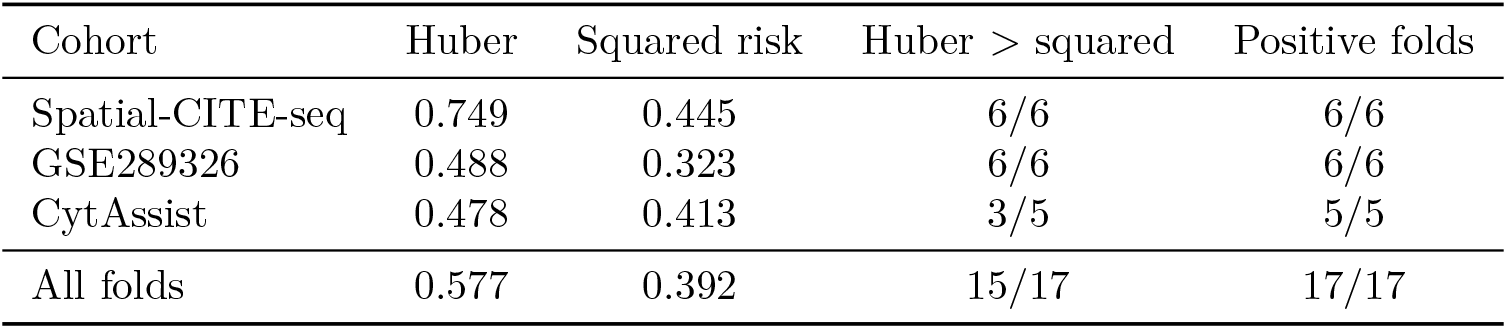
Huber-aligned and squared-risk value–mismatch diagnostics. Entries are mean within-fold Spearman associations with protein-level broad-minus-restricted gains.

| Cohort | Huber | Squared risk | Huber > squared | Positive folds |
|---|---|---|---|---|
| Spatial-CITE-seq | 0.749 | 0.445 | 6/6 | 6/6 |
| GSE289326 | 0.488 | 0.323 | 6/6 | 6/6 |
| CytAssist | 0.478 | 0.413 | 3/5 | 5/5 |
| All folds | 0.577 | 0.392 | 15/17 | 17/17 |

## 5 Conclusion

We studied spatial protein prediction through protein-specific center signals and broader molecular context. The value–mismatch characterization separates broad-context transfer into target-side value and source-to-target mismatch; for convex objectives, mismatch is related to cross-environment score variation, while squared loss gives an explicit covariance geometry. TSNR combines protein-specific center signals, broad molecular context, and bounded-score Huber regression.

TSNR achieves higher mean predictive performance than the reproduced baselines on the primary biological-holdout benchmark. Controlled analyses support both broader context and robust regression. Across the two external cohorts, contextual gains are positively associated with the Huber-aligned value–mismatch balance in all held-out folds; across all 17 holdouts, this diagnostic has higher mean association than the squared-risk specialization and is stronger in 15 folds. The association persists under spatially disjoint target blocks, while transfer-mismatch variation is more closely associated with protein-level contextual gain than target-side value alone. Together, these results suggest that contextual molecular scope and transfer robustness should be considered jointly rather than as independent design choices.

## A Theoretical Derivations

Section 3.3 characterizes transfer at the level of the complete signal–context predictor. This appendix gives the corresponding derivations, a restricted-curvature refinement of the transfer-mismatch bound, and the closed-form specialization under squared risk. Throughout, *s* denotes the pooled source distribution and *t* the held-out target environment. The representation maps are treated as fixed after being estimated from the source environments.

### A.1 Value–Mismatch Identity

For a fixed protein *p* and contextual regime *R* ∈ {*T, B*}, recall the prediction objective

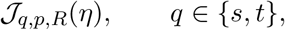

and let

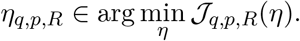

The source-trained broad-context advantage on the target environment is

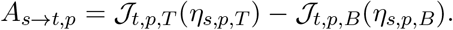

The target-side value difference is

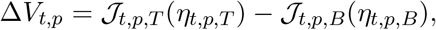

and the transfer mismatch under regime *R* is

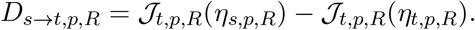

Because *η*_*t,p,R*_ minimizes the target objective,

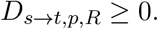

Define

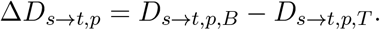

Expanding the definitions gives

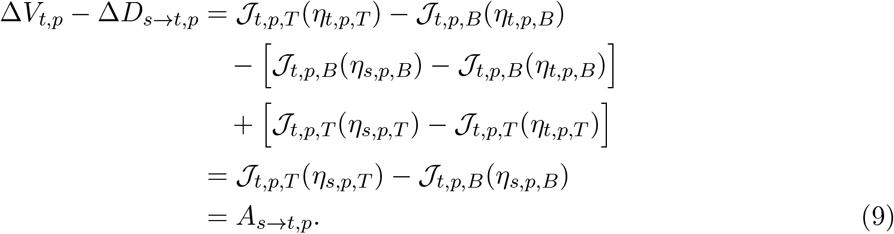

Therefore,

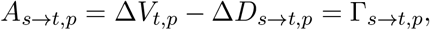

which establishes Eq. 3. Because the broad and target-restricted representations need not form nested predictor classes, Δ*V*_*t,p*_ and Δ*D*_*s*→*t,p*_ may each have either sign.

### A.2 Transfer Mismatch and Cross-Environment Score Shift

We next characterize the regime-specific transfer mismatch. To simplify notation, fix *p* and *R* and write

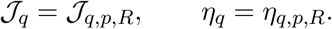

Suppose *J*_*t*_ is differentiable and convex in *η*. First-order convexity at *η*_*s*_ gives

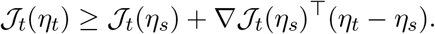

Rearranging yields

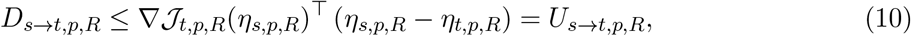

which proves Eq. 4. By Cauchy–Schwarz,

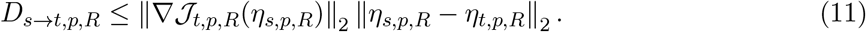

If *η*_*s,p,R*_ is an interior stationary source solution, then

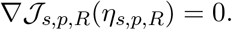

Hence

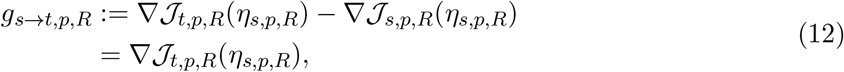

so the target score in Eq. 10 is precisely the source-to-target change in the estimating score evaluated at the source solution.

Combining Eq. 10 with the value–mismatch identity gives

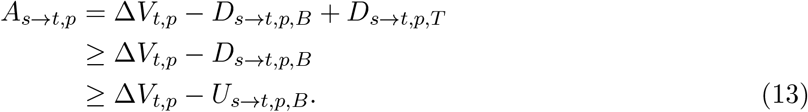

Therefore,

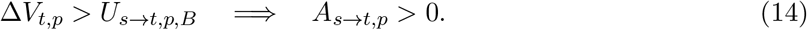

This is a sufficient, rather than necessary, condition for favorable transfer of the broader contextual representation.

### A.3 Restricted-Curvature Refinement

The first-order result above does not require strong convexity. A sharper norm-based bound follows when the target objective has positive curvature along the displacement from the source to the target solution.

For a fixed *p* and *R*, suppose that there exists *µ*_*p,R*_ *>* 0 such that

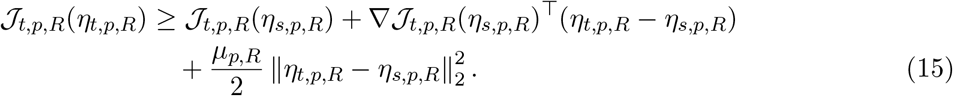

This condition is implied, for example, by *µ*_*p,R*_-strong convexity on a convex region containing the two solutions, but it only requires curvature along the relevant source–target displacement.

Rearranging Eq. 15 gives

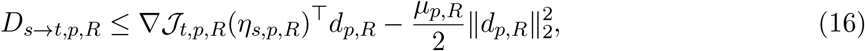

where

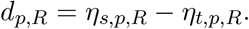

For any vectors *g, d* and *µ >* 0,

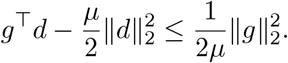

Applying this inequality to Eq. 16 yields

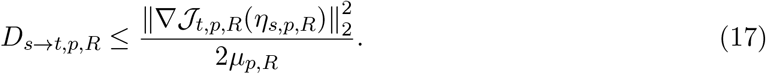

When the source solution is stationary, Eq. 12 gives the equivalent score-shift form

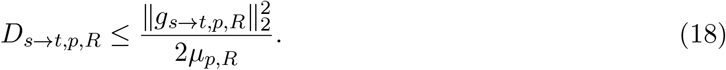

Consequently, a curvature-based sufficient condition for favorable broad-context transfer is

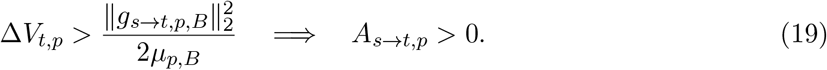

This refinement is local to the relevant predictor displacement and does not require the complete prediction objective to be globally strongly convex.

For a nonzero displacement, the largest pairwise curvature constant satisfying Eq. 15 with equality is

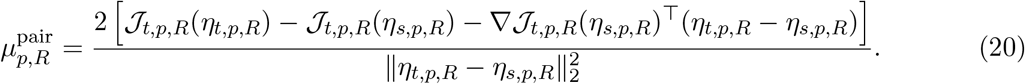

Convexity implies 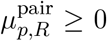. This quantity is useful for diagnosing curvature along the particular source–target displacement without making a global curvature assumption.

### A.4 Squared-Risk Geometry

We finally specialize the value–mismatch characterization to unregularized squared risk with a linear predictor. Let

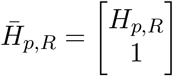

denote the complete signal–context representation augmented with an intercept, and let *w* be its coefficient vector. For *q* ∈ {*s, t*},

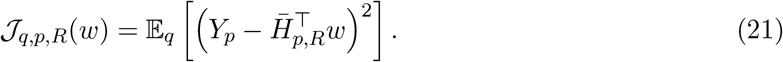

Define

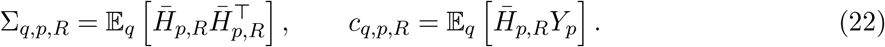

Expanding Eq. 21,

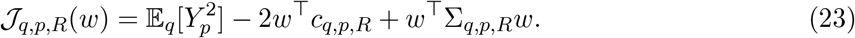

If Σ_*q,p,R*_ is nonsingular, the optimum is

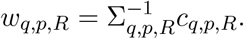

More generally, the canonical minimum-norm solution is

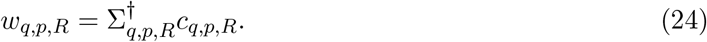

To see that this solution is well defined, let *v* lie in the null space of Σ_*q,p,R*_. Then 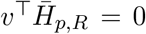 almost surely, and therefore *v*^⊤^*c*_*q,p,R*_ = 0. Hence *c*_*q,p,R*_ lies in the range of Σ_*q,p,R*_.

Using *c*_*t,p,R*_ = Σ_*t,p,R*_*w*_*t,p,R*_ in Eq. 23 gives, for any *w*,

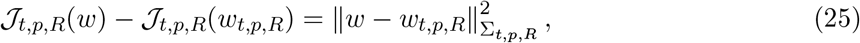

where 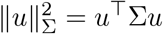. Evaluating this identity at the source solution yields

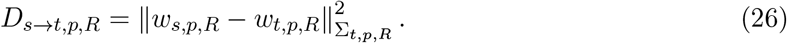

The target-optimal objective is

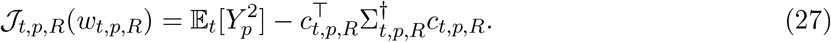

Therefore the target-side value difference between broad and target-restricted context is

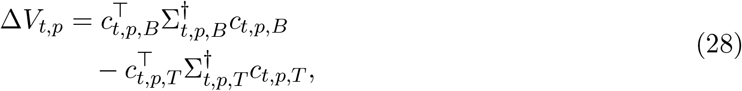

completing the squared-risk specialization described around Eq. 5.

The squared-risk case also gives an exact connection between transfer mismatch and score shift.

The target gradient at the source solution is

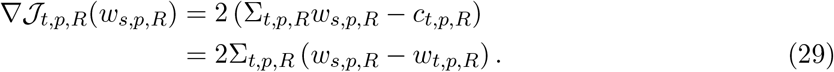

Since the squared-risk source optimum is stationary, this gradient is also the cross-environment score shift *g*_*s*→*t,p,R*_. Consequently,

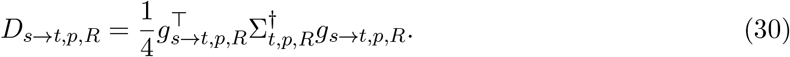

Thus, under squared risk, the general score-shift mechanism has an exact covariance-weighted geometry: mismatch is weighted by the pseudoinverse target covariance 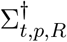, so score variation along lower-variance supported target directions contributes more strongly. Moreover,

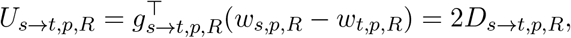

which makes the relation between the general first-order bound and the quadratic specialization explicit.

## B Experimental and Reproducibility Details

### B.1 Development and Configuration Selection

The six-section Spatial-CITE-seq benchmark served as the primary evaluation benchmark. Within every leave-one-section-out fold, the held-out section is excluded from gene ranking, context projection, regression fitting, and all other fold-dependent transformations. The reported TSNR configuration was fixed before the sensitivity analyses in Appendix C.4.

The finalized TSNR configuration was subsequently applied unchanged to GSE289326 and CytAssist. Cohort-specific assay preprocessing and transcriptomic input spaces are described in Appendix B.2. Baseline implementations and protocol-specific adaptations are described in Appendix B.4.

### B.2 TSNR Implementation

All fold-dependent transformations are fitted using outer-training data only. For each protein, TSNR selects *K* = 256 center genes using the cross-environment association score in Eq. 6. Pearson associations are computed separately within each source environment after the cohort-specific preprocessing described below, and the selected gene set is recomputed for every outer fold.

For the six-section Spatial-CITE-seq benchmark, we use the processed post-QC in-tissue spots. RNA counts are normalized to a total count of 10^4^ per spot, log1p-transformed, and gene-wise scaled with values clipped at 10. The 11,535-gene input space is the fixed intersection of gene names across the six sections.

For GSE289326 and the CytAssist cohort, RNA is normalized to 10^4^ counts per spot and log1p-transformed, followed by per-section gene-wise standardization and clipping to [−10, 10]. These cohorts use their 18,085-gene transcriptomic input spaces. This cohort-specific RNA preprocessing is applied independently within each section and does not use protein responses. All subsequent fold-dependent predictor and response transformations are estimated from source observations only.

Protein responses follow the finalized cohort-specific assay representation: the primary Spatial-CITE-seq benchmark uses centered log-ratio (CLR)-transformed protein measurements, whereas GSE289326 and CytAssist retain their native protein count representation. The 31 assay features correspond to 30 unique protein identities because PTPRC is represented by two antibody features, PTPRC_1 and PTPRC_2. Within every outer fold, the corresponding source-derived fold-dependent transformations are applied to the held-out environment where applicable.

Spatial neighborhoods are constructed independently within each tissue section using Euclidean distances between spot coordinates. We construct a 36-nearest-neighbor candidate graph with the center spot excluded and use neighbor ranks 1–6 to define the local neighborhood. Neighborhood profiles are formed in the cohort-specific transcriptomic space described above.

The PCA context transform is fitted using neighborhood profiles from the outer-training environments only, and the first *d*_*z*_ = 64 components are retained. Held-out neighborhood profiles are transformed using this training-derived basis without refitting.

A separate HuberRegressor from scikit-learn 1.9.0 (Pedregosa et al., 2011) is fitted for each protein with *ϵ* = 1.35, *α* = 10^−4^, `max_iter`=500, `fit_intercept`=True, tol=10^−5^, and `warm_start`=False. The estimator jointly fits the regression coefficients, intercept, and residual scale; the *ℓ*_2_ penalty with strength *α* is applied to the coefficients but not the intercept. No sample weights are used. The same estimator configuration is used across all 17 biological holdout folds.

### B.3 Controlled Ablation Construction

The controlled variants keep the protein-specific center representation and final regression model fixed and modify only the contextual branch. The *center-only* variant removes the contextual branch entirely. The *target-restricted* variant uses the same protein-specific gene set *G*_*p*_ as the center representation, averages these genes over the same rank-1–6 spatial neighbors as TSNR, and fits a protein-specific 64-dimensional PCA context transform using outer-training data only. TSNR instead constructs the neighborhood profile from the full cohort-specific transcriptomic input space before applying the shared training-only PCA64 transform.

#### Broad context excluding *G*_*p*_

As an additional molecular-scope control, we remove each protein’s selected center-gene set *G*_*p*_ from the broad contextual transcriptomic input while retaining the center representation and the same outer biological holdout protocol. The resulting broad-minus-restricted gain is +0.0131, compared with +0.0150 for the full broad context; it is positive in 4/6 biological holdouts, and the aggregate score is only 0.0018 below full broad context.

The *correspondence-disrupted* control tests center–context correspondence. The broad PCA64 context representation is first constructed exactly as in TSNR. The resulting context vectors are then randomly permuted across spots within each biological section before the protein-specific regressor is refitted. The PCA basis is therefore fitted before the permutation and is not refitted after correspondence is disrupted. The permutation is not constrained to be a derangement, so occasional self-matches are possible. On the primary benchmark, we average three fixed permutations generated with seeds 142–144.

The *random-gene* control tests molecular content while preserving the spatial neighborhood. For each protein and random seed, 256 genes are sampled without replacement from the available transcriptomic gene pool after excluding that protein’s selected center genes. Their rank-1–6 neighborhood averages are compressed with the same protein-specific PCA64 construction used in the target-restricted variant, and the same downstream protein-specific regressor is fitted. Random sets are drawn independently for each protein and seed. The reported primary-benchmark result averages five fixed draws generated with seeds 42–46.

### B.4 Baseline Implementations

#### DGAT

We use the official DGAT implementation under the same six outer leave-one-section-out splits as TSNR. DGAT receives the complete set of 11,535 shared genes. Its graph construction, model architecture, training objectives, and optimization procedure follow the official implementation. Protein measurements from the held-out section are not used for training or model selection.

#### SR2P

We retain the official cross-sample feature-selection and stacking procedure. Within each outer fold, highly variable genes are selected within the training samples and intersected across samples according to the original cross-sample workflow. Under the six outer folds used here, this procedure yields 459–514 genes per fold. Predictions used to train the stacking model are generated by leave-one-training-section-out cross-validation, so the outer held-out section is not used to construct meta-learner training features.

#### SPINE

SPINE is evaluated with the same outer biological partitions and the complete 11,535-gene input. Model selection is performed using a validation section contained within the outer-training set. Spatial graph operations are restricted to individual biological sections. To use the same protein target representation and evaluator across methods, we use the CLR target transformation supported by the official SPINE preprocessing utilities. This differs from the row-wise min–max transformation used in its primary reported setting.

## C Additional Analyses

### C.1 Huber Value–Mismatch Diagnostic

Section 4.4 evaluates the value–mismatch characterization using the same scale-adaptive Huber estimator used by TSNR. For each biological holdout *t*, protein *p*, and contextual regime *R* ∈ {*T, B*}, the center and context representations are constructed exactly as in the corresponding source-trained prediction model and are then treated as fixed.

Let

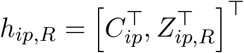

denote the complete representation for observation *i*. For parameters *η* = (*w, b, σ*) and residual

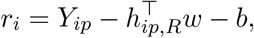

the finite-sample objective corresponding to the scikit-learn 1.9.0 HuberRegressor implementation used in the experiments is

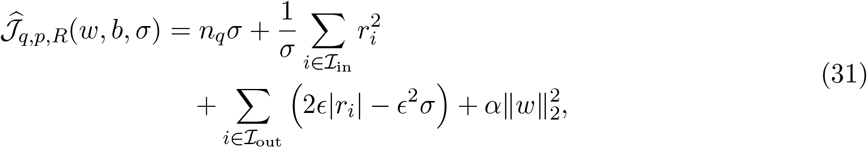

where

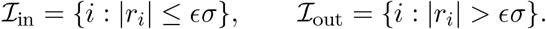

The experiments use *ϵ* = 1.35 and *α* = 10^−4^, as specified in Appendix B.2. The intercept and residual scale are fitted jointly with the coefficient vector, while the *ℓ*_2_ penalty is applied only to the coefficients.

For each protein and regime, 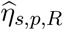 denotes the predictor fitted on the pooled source environments. We additionally obtain a target-reference diagnostic fit 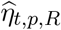 using the same objective on the held-out environment while keeping the source-derived representation fixed. These target-side fits are used only for the analysis in Sec. 4.4.

The empirical transfer mismatch is

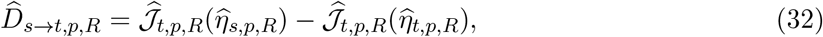

and the target-side value difference is

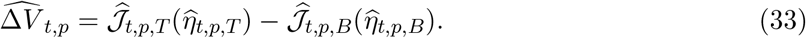

We then compute

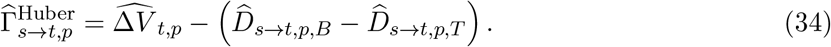

The target-side fits and protein measurements above are used only for this retrospective characterization. The reported TSNR predictions continue to use the source-trained predictors 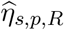.

### C.2 Additional Value–Mismatch Results

#### Value and mismatch components

To examine which part of the decomposition carries the protein-level ordering, we compare the complete balance with its two components. For comparability of objective magnitudes across cohorts, this analysis uses the normalized empirical objective 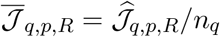. Within a biological fold, this is a positive rescaling and therefore does not alter the rank ordering of 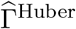.

For CytAssist, the normalized target-side value difference is positive for 96% of protein–fold pairs (mean Δ*V* ≈ +0.037), but its mean mismatch increase is larger (Δ*D* ≈ +0.052), yielding mean Γ ≈ −0.015.

#### Spatially disjoint target validation

We further separate the target measurements used to construct the diagnostic from those used to evaluate predictive gain. Within each held-out tissue, spatial coordinates are standardized and split at the median of their first principal component. We estimate the Huber value–mismatch diagnostic on one spatial block and compute broad-minus-restricted predictive gain on the other, then reverse the two blocks.

Across all 34 directional comparisons, the mean Spearman correlation is 0.480. Averaging the two directions within each biological holdout gives a positive association in all 17 folds.

#### Comparison with the squared-risk specialization

The closed-form squared-risk geometry in Sec. 3.3 provides a complementary analytic specialization of the same value–mismatch framework. We therefore also compute the squared-risk diagnostic using the complete signal–context representation.

### C.3 Empirical Diagnostics for the Score-Shift Bound

We evaluate the first-order mismatch bound from Eq. 4 using the normalized empirical objective

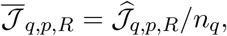

which preserves each fitted minimizer. Define

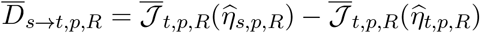

and

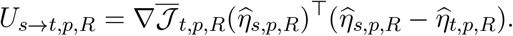

Across the 1,054 protein–fold–regime fits, *D* and *U* have a Spearman correlation of approximately 0.98, with median *D/U* = 0.648.

### C.4 Predictor and Representation Sensitivity

#### Predictor and compact-representation sensitivity

We evaluate alternative regression estimators and compact context representations under the same six outer biological holdout splits and evaluation metric. Table 8 contains the full set of estimator–representation combinations used in this sensitivity analysis.

**Table 8:** Predictor and context-representation sensitivity on the six-section Spatial-CITE-seq benchmark. The reported TSNR configuration is PCA64 with Huber regression; FA64 denotes a 64-dimensional factor-analysis representation implemented with scikit-learn (Pedregosa et al., 2011).

| Context representation | Regression estimator | Mean macro Spearman |
|---|---|---|
| PCA64 | Ridge | 0.4320 |
| PCA64 | Huber | 0.4391 |
| PCA64 | Elastic Net | 0.4398 |
| PCA64 | HistGradientBoosting | 0.4404 |
| PCA64 | MLP | 0.4228 |
| FA64 | Ridge | 0.4345 |
| FA64 | Huber | 0.4410 |
| FA64 | Elastic Net | 0.4399 |
| FA64 | HistGradientBoosting | 0.4394 |
| PCA256 | Ridge | 0.4367 |
| PCA256 | Huber | 0.4408 |

**Table 9:** Sensitivity to the number of protein-specific center genes on the six-section Spatial-CITE-seq benchmark. All other TSNR components are fixed to the reported configuration.

| *K* | Tonsil-1 | Tonsil-2 | Breast | GBM | Meso-1 | Meso-2 | Mean |
|---|---|---|---|---|---|---|---|
| 64 | .5072 | .6575 | .5812 | .2007 | .3332 | .3364 | .4360 |
| 128 | .5112 | .6570 | .5865 | .2034 | .3371 | .3410 | .4394 |
| 256 | .5142 | .6573 | .5899 | .1971 | .3346 | .3414 | .4391 |
| 512 | .5206 | .6538 | .5872 | .2069 | .3337 | .3369 | .4399 |
| 1024 | .5249 | .6485 | .5701 | .2056 | .3251 | .3236 | .4330 |

**Table 10:** Controlled comparison of compact context representations under the same protein-specific center representation and Ridge regression estimator. AE64 denotes a learned 64-dimensional autoencoder bottleneck.

| Context representation | Mean macro Spearman |
|---|---|
| PCA64 | 0.4320 |
| AE64 | 0.4212 |
| NMF64 | 0.4356 |

Performance is similar across most estimator–representation combinations, indicating that the representation-level conclusions are not concentrated in a single compact projection or regression estimator.

We additionally screened a frozen pretrained spatial-transcriptomics representation under the same primary-benchmark Ridge protocol. It did not improve over the corresponding PCA64 context baseline.

#### Sensitivity to center-gene set size

We vary the number of protein-specific center genes while keeping the remaining TSNR configuration fixed.

Performance varies by only 0.0069 in mean macro Spearman across *K* = 64 to *K* = 1024, with no monotonic trend.

#### Learned and alternative context representations

We also compare PCA with a learned autoencoder and NMF implemented using scikit-learn (Pedregosa et al., 2011), while fixing the protein-specific center representation, the outer biological splits, the evaluation metric, and Ridge regression.

The autoencoder uses the same preprocessed neighborhood-mean transcriptomic profiles as PCA64 and a 11,535 → 512 → 128 → 64 encoder with a symmetric decoder. It is trained with mean-squared reconstruction loss using AdamW (Loshchilov and Hutter, 2019) with learning rate 10^−3^, weight decay 10^−4^, batch size 2048, and early stopping on a section held out from the outer-training environments.

### C.5 Additional Robustness Analyses

#### Breadth of the primary-benchmark improvement

Across the 186 protein–fold TSNR– DGAT differences, 65.1% are positive, with median Δ*ρ* = +0.0195. After averaging within protein across the six biological folds, 20/31 proteins have positive differences, with median +0.0245. At the biological-fold level, TSNR is higher than DGAT in 5/6 holdouts.

#### Huber versus Ridge

The main-text comparisons use the same signal–context representation for Huber and Ridge. Penalty-matched Ridge uses *α* = 10^−4^. For the source-tuned Ridge comparator, we select regularization using source-only leave-one-source-environment-out validation over

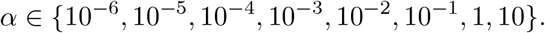

#### Residual behavior

We characterize residual behavior using out-of-fold predictions from inner leave-one-training-environment-out Ridge fits on the primary benchmark. Across the 186 protein– fold residual groups, the median excess kurtosis is 2.33 and all 186 groups have positive excess kurtosis. Approximately 2.7% of standardized residuals exceed three absolute standard deviations and 0.3% exceed five. The largest 1% of residuals account for approximately 14.8% of the total squared loss.

These residuals exhibit moderate heavy-tailed behavior and concentration of squared loss in a small fraction of observations, consistent with the use of a regression loss whose residual score is bounded.

#### Response-contamination stress test

Using Ridge with fixed regularization (*α* = 1.0), we perturb source-training responses over contamination fractions *c* ∈ {0, 0.01, 0.02, 0.05}, with perturbation magnitude scaled by the median absolute deviation of the response. At *c* = 0.05 and *κ* = 8 MAD(*y*), mean macro Spearman decreases from 0.4319 to 0.4254 for Ridge (−0.0065) and from 0.4391 to 0.4386 for Huber (−0.0005).

## Notes

https://www.10xgenomics.com/datasets/gene-protein-expression-library-of-human-tonsil-cytassist-ffpe-2-standard

https://www.10xgenomics.com/datasets/visium-cytassist-gene-and-protein-expression-library-of-human-tonsil-with-add-on-antibodies-h-e-6-5-mm-ffpe-2-standard

https://www.10xgenomics.com/datasets/gene-and-protein-expression-library-of-human-breast-cancer-cytassist-ffpe-2-standard

https://www.10xgenomics.com/datasets/gene-and-protein-expression-library-of-human-glioblastoma-cytassist-ffpe-2-standard

https://www.ncbi.nlm.nih.gov/geo/query/acc.cgi?acc=GSE300851

https://www.ncbi.nlm.nih.gov/geo/query/acc.cgi?acc=GSE289326

https://doi.org/10.5281/zenodo.17999961

### Competing Interest Statement

The authors have declared no competing interest.

